# Formal Links between Feature Diversity and Phylogenetic Diversity

**DOI:** 10.1101/2020.04.06.027953

**Authors:** Kristina Wicke, Arne Mooers, Mike Steel

**Affiliations:** Institute of Mathematics and Computer Science, University of Greifswald, Greifswald, Germany; Department of Biological Sciences and the Crawford Lab for Evolutionary Studies, Simon Fraser University, Burnaby, British Columbia, V5A1S6, Canada; Biomathematics Research Centre, University of Canterbury, Christchurch, New Zealand

**Keywords:** Phylogenetic diversity, feature diversity, evolutionary distinctiveness, Shapley value

## Abstract

The extent to which phylogenetic diversity (PD) captures feature diversity (FD) is a topical and controversial question in biodiversity conservation. In this short paper, we formalise this question and establish a precise mathematical condition for FD (based on discrete characters) to coincide with PD. In this way, we make explicit the two main reasons why the two diversity measures might disagree for given data; namely, the presence of certain patterns of feature evolution and loss, and using temporal branch lengths for PD in settings that may not be appropriate (e.g. due to rapid evolution of certain features over short periods of time). Our paper also explores the relationship between the ‘Fair Proportion’ index of PD and a simple index of FD (both of which correspond to Shapley values in cooperative game theory). In a second mathematical result, we show that the two indices can take identical values for any phylogenetic tree, provided the branch lengths in the tree are chosen appropriately.

## Introduction

Almost 30 years ago, Dan Faith published a seminal paper that laid out how phylogenies might aid in identifying sets of species with maximal “feature diversity” (Faith, 1992). Faith’s stated goal was to support practical biodiversity conservation in the face of limited resources, coupled with the assumption that maximising feature diversity (the total number of unique character states represented by a set of taxa) was a desirable conservation target.

Drawing on the call of Vane-Wright et al. (1991) to consider taxonomic distinctiveness when prioritizing species, Faith introduced the phylogenetic diversity (PD) metric, simply the sum of the edge lengths of the minimal subtree linking a subset of species to the root of the encompassing phylogeny (also called the ‘minimum spanning path’ by Faith (1992)). Importantly, these edge lengths were given in units of reconstructed character changes under maximum parsimony on the cladogram representing a character state matrix with no homoplasy. Faith showed, with an example, that the sum of these reconstructed edge lengths would lead to the same total feature diversity as that calculated from the character matrix itself. Importantly, if these cladistic edge lengths are representative of all features, then maximising PD (e.g. over a given subset size) would maximise feature diversity, even in the face of some homoplasy. The bulk of Faith’s 1992 paper was devoted to introducing the machinery to maximise PD.

Efficient algorithms for finding maximum PD sets are available (Bordewich et al. (2008)), the metric has been extended to networks (Minh et al. (2009)), and there are countless case studies that both measure and optimize PD for conservation (see, e.g., Pollock et al. (2017)); Faith’s original paper has been cited in excess of 2000 times. A recent review (Tucker et al., 2019) considered the literature concerning both the empirical correlations between PD and feature diversity, and the expected relationship between PD and various conservation values.

Surprisingly, though, the necessary conditions under which PD will capture feature diversity have never been formalized. Here, by using discrete characters, a model with no homoplasy, and appropriate edge lengths, we prove that the PD of a subtree does indeed measure feature diversity as defined by (Faith, 1992). This proof allows us to state more formally when PD does not necessarily capture feature diversity, thereby allowing for further modelling and statistical evaluation of the expected relationship under more realistic models. Given the close connection between PD and taxonomic distinctiveness, we also consider the conditions under which its phylogenetic measure (specifically the Shapley value of evolutionary isolation) can capture its feature-based analogue.

## Preliminaries

### Feature diversity

Consider a set *X* of taxa with |*X*| = *n*, and suppose that each taxon *x* ∈ *X* has an associated finite set *F*_*x*_ of ‘features’. To allow extra generality, we will assume that each element *f* ∈ *F*_*x*_ has a corresponding positive score *µ*(*f*) ∈ ℝ^>0^, which might be viewed as a measure of the complexity, novelty, or richness of *f* (the default option is to set *µ*(*f*) = 1 for all *f*). Let ℱ denote the set of all features present amongst the taxa in the collection *X*, and let 𝔽 = (*F*_*x*_ : *x* ∈ *X*) be the ordered *n*-tuple containing the feature sets of the taxa in *X*. We will sometimes call 𝔽 a *feature assignment* as it summarizes how a set of features is assigned to each taxon in *X*.

Note that 𝔽 provides the same information as a table showing the presence and absence of features across taxa. So if *X* = {*a, b, c*}, then the feature assignment 𝔽 = (*F*_*a*_, *F*_*b*_, *F*_*c*_) where *F*_*a*_ = {*f*_1_, *f*_2_}, *F*_*b*_ = {*f*_1_, *f*_3_} and *F*_*c*_ = {*f*_2_, *f*_3_}, corresponds to a standard character state matrix where there are two states per feature: presence (1) or absence (0) (see Table 1).

**Table 1.** *A standard character state matrix (*0 =*absence*, 1 =*presence) representing the assignment of three features (f*_1_, *f*_2_, *f*_3_*) across three taxa (a, b, c*).

| Taxon | $f_1$ | $f_2$ | $f_3$ |
| --- | --- | --- | --- |
| $a$ | 1 | 1 | 0 |
| $b$ | 1 | 0 | 1 |
| $c$ | 0 | 1 | 1 |

Given a subset *Y* of *X*, let

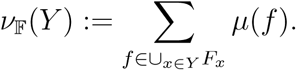

Thus, *ν*_𝔽_(*Y*) is the sum of the values of the features that are present in at least one taxon in *Y*. We refer to *ν*_𝔽_(*Y*) as the *feature diversity* (FD) of *Y*. Note that in this sum, each feature is counted only once if present.

The function *ν*_𝔽_ (which assigns each subset *Y* of *X* a non-negative real value *ν*_𝔽_(*Y*)) clearly satisfies the following two properties: *ν*_𝔽_(ø) = 0 and *ν*_𝔽_ is monotone (i.e. *Y* ⊆ *Y*′ ⇒ *ν*_𝔽_(*Y*) ⩽ *ν*_𝔽_(*Y*′)). Moreover, *ν*_𝔽_ also satisfies the submodularity inequality:

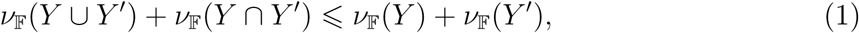

and a proof is provided in the ‘Proofs of Propositions’ section.

### Phylogenetic diversity

Now consider a rooted phylogenetic *X*-tree *T* = (*V, E*) with root *ρ*, leaf set *X*, and edge length assignment ℓ : *E* → ℝ^⩾0^. For technical reasons (by allowing greater generality in the statement of our results) we assume that *T* has an additional ‘stem edge’ (*ρ*′, *ρ*) where *ρ*′ is a degree-1 vertex and *ρ* has in-degree 1 and out-degree at least 2 (see Fig. 1). The *phylogenetic diversity* (PD) of a subset *Y* of *X* is usually defined as the sum of the lengths of the edges in the minimal subtree of *T* that contains the leaves in *Y* and the root *ρ* of *T*. Here, we extend this definition by also including the length of the stem edge (*ρ*′, *ρ*) in the calculation of PD for any subset *Y* ⊆ *X* with |*Y* | ⩾ 1. This adds a constant, namely ℓ((*ρ*′, *ρ*)), to all subsets *Y* ⊆ *X* \ ø but does not affect properties of PD, such as its monotonicity or submodularity.

**Fig. 1.**
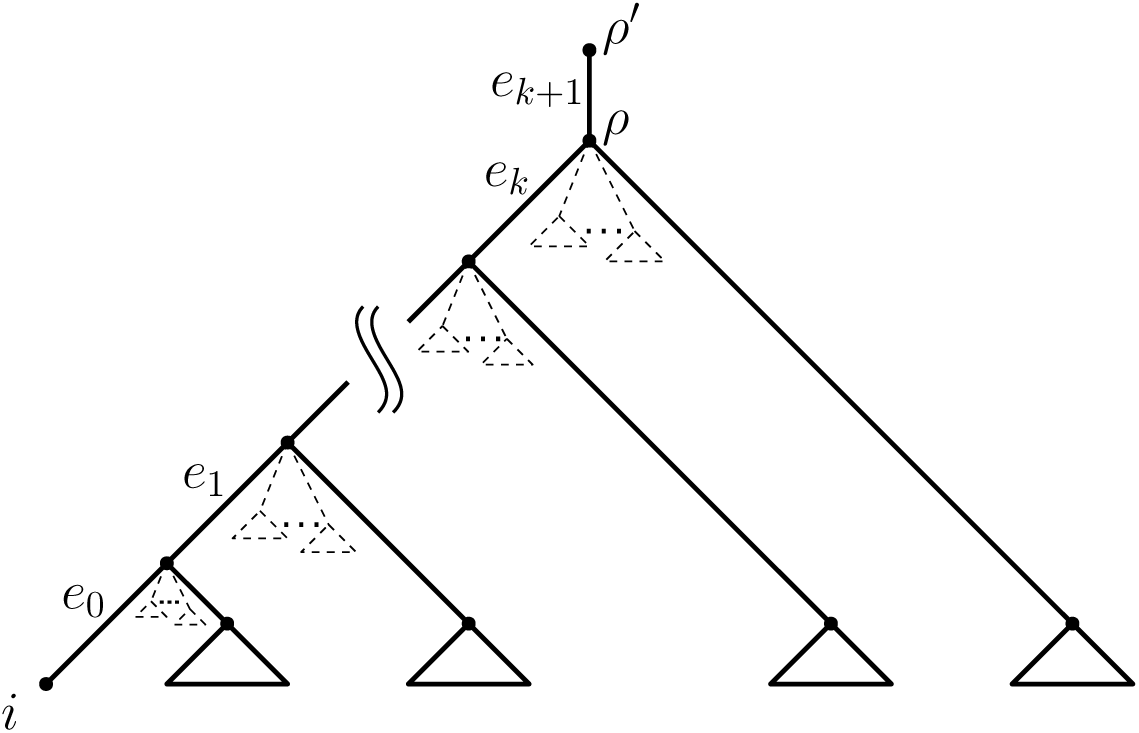
Representing a phylogenetic *X*-tree *T* relative to a reference leaf *i*. Note that *T* is not assumed to be binary.

**Fig. 2.**
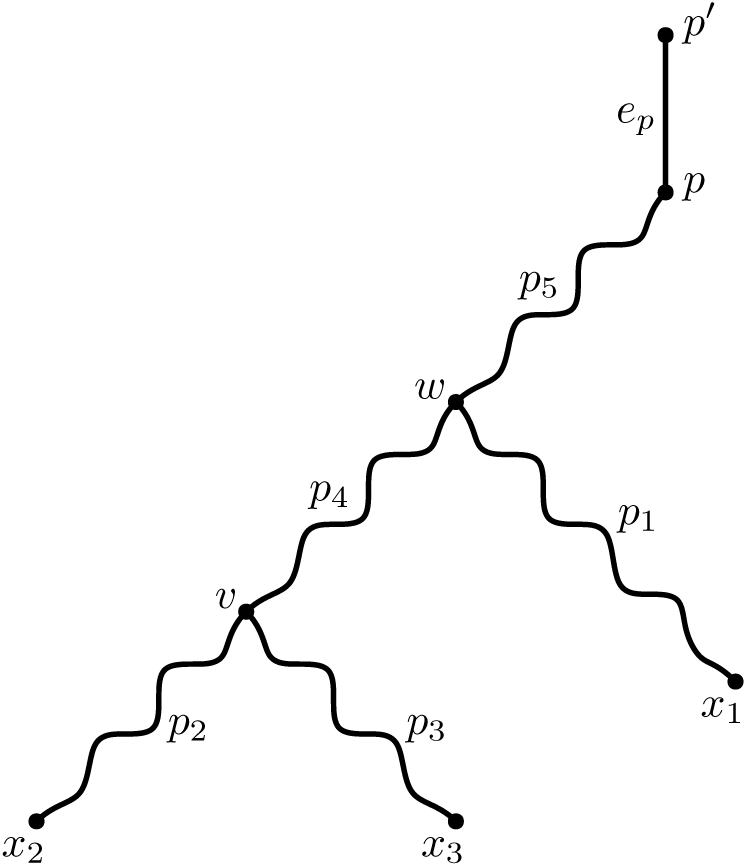
Subtree induced by taxa *x*_1_, *x*_2_, and *x*_3_ in the proof of Lemma 2. *v* denotes the most recent common ancestor of *x*_2_ and *x*_3_. Analogously, *w* denotes the most recent common ancestor of *x*_1_, *x*_2_, and *x*_3_. Furthermore, *p*_1_ denotes the unique path from *w* to *x*_1_, *p*_2_ denotes the unique path from *v* to *x*_2_ and so forth.

## Linking feature diversity to phylogenetic diversity

Next consider a model, based on a rooted phylogenetic *X*–tree *T* in which (i) each feature in ℱ arises on exactly one edge of *T* and (ii) each feature that arises is never lost and is passed down to all descendant vertices (including the leaves). This is just a model where every feature is a perfect synapomorphy.

We can describe this more precisely by specifying a map *h* : ℱ → *E*, which indicates which edge each given feature arises on (note that several features may arise on the same edge). Thus *h*^−1^(*e*) denotes the set of features that arise on edge *e*. Here, we assume that *h*^−1^(*e*) ≠ ø for all interior edges of *T* (i.e., each interior edge of *T* gives rise to at least one feature). Notice that this is equivalent to allowing interior edges with ‘no event’ (i.e., without a feature arising on them) and then contracting all interior ‘no event’ edges.

Note, however, that there may be pendant edges incident to leaves of *T*, on which no features arise. Furthermore, we also consider the stem edge (*ρ*′, *ρ*) of *T* to be a pendant edge; in particular, no features arise on this stem edge precisely when there is no feature that is present in every taxon.

Under this model, *F*_*x*_ is then equal to the union of the sets *h*^−1^(*e*) over all the edges *e* on the (unique) path from *ρ*′ to leaf *x*.

When a feature assignment 𝔽 can be realized in this way, we will denote this by writing 𝔽 = *F* [*T, h*]. Not every feature assignment 𝔽 can be realized in this way (on any tree). As an example, consider the feature assignment described by the character state matrix above (Table 1). In this case, there is no rooted phylogenetic *X*–tree *T* = (*V, E*) and map *h* : {*f*_1_, *f*_2_, *f*_3_} → *E* for which (*F*_*a*_, *F*_*b*_, *F*_*c*_) = *F* [*T, h*].

Fortunately, it is easy to characterise precisely when a feature assignment 𝔽 can be realized as *F*[*T, h*], and where *T* is either stipulated or not. The required condition corresponds to the well-known structure of characters necessary (and sufficient) to perfectly fit a common phylogenetic tree, namely that character states are arranged among taxa as a set of nested apomorphies.

To describe this, we first introduce some additional notation. Let *X*_*f*_ := {*x* ∈ *X* : *f* ∈ *F*_*x*_} denote the subset of taxa in *X* that have feature *f*. Moreover, let 𝒞_ℱ_ := {*X*_*f*_ : *f* ∈ ℱ} be the collection of the sets *X*_*f*_. The following result easily follows from other well-known results in phylogenetics; however, for completeness a self-contained proof is given in the ‘Proofs of Propositions’ section.

### Proposition 1

i. 𝔽 = *F*[*T, h*] for some *h* : ℱ → *E* if and only if *X*_*f*_ corresponds to a cluster of *T* for each feature *f* ∈ ℱ. Moreover, when 𝔽 = *F*[*T, h*], the map *h* is uniquely determined: for each *f* ∈ ℱ, *h*(*f*) is the edge directly above the most recent common ancestor of the taxa in *X*_*f*_.
ii. There exists a tree *T* and map *h* such that 𝔽 = *F*[*T, h*] if and only if 𝒞_ℱ_ is a hierarchy on *X*. In other words, for all pairs *X*_*f*_, *X*_*f*′_ ∈ 𝒞_ℱ_, we have *X*_*f*_ *∩ X*_*f*′_ ∈ {ø, *X*_*f*_, *X*_*f*′_} (i.e. *X*_*f*_ and *X*_*f*′_ are either disjoint or nested).

## First main result

We can now describe the relationship between FD and PD in a precise way.

### Theorem 1

Let *T* be a rooted phylogenetic *X*–tree and let 𝔽 be an assignment of features across the taxa in *X*.

i. 𝔽 = *F*[*T, h*] for some function *h* : ℱ → *E* if and only if *ν*_𝔽_ is exactly equal to the PD function for some edge length assignment ℓ of *T* that assigns strictly positive lengths to all interior edges of *T* and non-negative lengths to all pendant edges and the stem edge (*ρ*′, *ρ*) (i.e. *ν*_𝔽_(*Y*) = PD_(*T*,ℓ)_(*Y*) for all subsets *Y* of *X*).
ii. When (i) holds, *h* and ℓ are both uniquely determined. In particular, ℓ = ℓ_*h*_, where, for each *f* ∈ ℱ, ℓ_*h*_(*e*) := ∑_*f* :*h*(*f*)=*e*_ *µ*(*f*) (and ℓ_*h*_(*e*) = 0 for each pendant edge *e* of *T* with *h*^−1^(*e*) = ø).

### Proof of Theorem 1.

The proof of Theorem 1 relies on three lemmas (whose proof is given in the Appendix). We state these lemmas now, and then use them to establish Theorem 1.

### Lemma 1

Given a rooted phylogenetic *X*–tree *T*, suppose that 𝔽 = *F*[*T, h*] for some map *h* : ℱ → *E*. Let ℓ_*h*_ : *E* → ℝ^⩾0^ be defined by setting

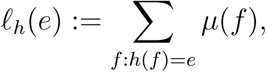

for each edge *e* of *T* (where ℓ_*h*_(*e*) := 0 if *h*^−1^(*e*) = ø). Then, for all subsets *Y* of *X* we have:

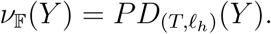

### Lemma 2

Given a rooted phylogenetic *X*-tree *T*, suppose that the identity

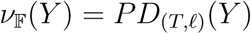

holds for all subsets *Y* ⊆ *X*, where ℓ : *E* → ℝ^⩾0^ is such that the interior edges of *T* are assigned strictly positive lengths and pendant edges (including the stem edge) are assigned non-negative lengths. Then, there exists a map *h* : ℱ → *E* such that 𝔽 = *F*[*T, h*].

### Lemma 3

Let *T* be a rooted phylogenetic *X*-tree (with additional stem edge). Then, the edge lengths of *T* are uniquely determined by the induced PD scores of all subsets *Y* ⊆ *X* with |*Y* | ⩽ 2.

We now show that Theorem 1 follows from these lemmas. Part (i) of Theorem 1, namely

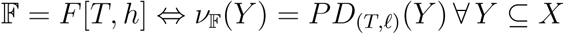

follows from Lemmas 1 and 2 (the ‘only if’ implication is implied by Lemma 1 and the ‘if’ implication is implied by Lemma 2).

For Part (ii), the uniqueness of ℓ (i.e., ℓ = ℓ_*h*_), follows by combining Lemmas 1 and 3. More precisely, Lemma 1 states that assigning edge lengths according to ℓ_*h*_ induces the equality of *ν*_𝔽_(*Y*) and 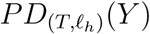 for all *Y* ⊆ *X*, whereas, by Lemma 3, the edge lengths of a given tree *T* are uniquely determined by the induced PD scores of all *Y* ⊆ *X* (indeed, even those with size at most 2 suffice). Moreover, the uniqueness of *h* is implied by Proposition 1, Part (i). This completes the proof. □

## Diversity indices

A *diversity index* for FD (or PD) is a score assigned to each taxon *x* ∈ *X* that sums to the total FD (or PD, respectively) of *X*. Diversity indices can be viewed as a way to apportion FD (or PD) fairly among the extant taxa. Although there are various ways to do this, we focus on one that is characterised by simple axioms, namely, the Shapley value (from cooperative game theory), which coincides, in the PD setting, with the well-known Fair Proportion index (described below).

### Feature diversity index

Given 𝔽, let

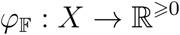

be the function defined by:

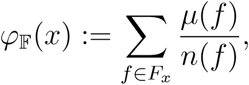

where *n*(*f*) is the number of taxa that have feature *f* (i.e. *n*(*f*) = |*X*_*f*_ |). In words, *φ*_𝔽_(*x*) assigns to each taxon *x* a sum of scores — one score for each of its features — where the score for feature *f* is *µ*(*f*) if *x* is the only taxon having this feature; otherwise, the score equals *µ*(*f*) divided by the total number of taxa having feature *f*.

The following result provides a formal justification for regarding *φ*_𝔽_ as a natural index of FD. Note that this index does not depend on any underlying phylogeny, or on assumptions concerning feature evolution, and is easily computed.

The result is phrased within the general framework of cooperative game theory (a topic more well-known in economics than biology, though it has recently been applied to PD, as we discuss below). In this general framework, one has a finite set *X* and a function *s* that assigns to each subset *Y* of *X* a corresponding score *s*(*Y*) with *s*(ø) = 0 (in our current setting *s*(*Y*) = *ν*_𝔽_(*Y*)). Given the pair (*X, s*), one seeks to apportion the score of the full set *X* among each of its elements according to an index (i.e., a value for each element of *X*) in a way that reflects the contribution each element makes to the total score. In this general framework, there is a particular index, called the *Shapley value*, that is uniquely determined by well-motivated axioms, and which is given by an explicit (if somewhat complex) combinatorial expression (Shapley, 1953).

#### Proposition 2

The FD index *φ*_𝔽_ is precisely the Shapley value for the pair (*X, ν*_𝔽_). In particular, Σ_*x*∈*X*_ *φ*_𝔽_(*x*) = *ν*_𝔽_(*X*).

The proof of Proposition 2 is provided in the ‘Proofs of Propositions’ section.

### Phylogenetic diversity index

Given the pair (*T*, ℓ), the *Fair Proportion index* (FP) (from Redding (2003) and Redding et al. (2007), see also Isaac et al. (2007)) for taxon *x* is given by:

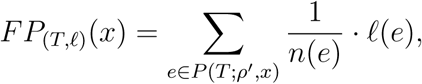

where *P* (*T*; *ρ*′, *x*) denotes the unique path from *ρ*′ to *x* and where *n*(*e*) is the number of leaves descending from the endpoint of edge *e* closest to the leaves.

It turns out that the FP index coincides exactly with the Shapley value based on PD (i.e. when PD is used as the characteristic function in the underlying cooperative game), a result first shown by Fuchs and Jin (2015). As *φ*_𝔽_ is (by Proposition 2) equivalent to the Shapley value based on FD, Theorem 1 thus has the interesting implication that if a feature assignment 𝔽 can be realized on a tree (i.e., if 𝔽 = *F*[*T, h*]), then the Shapley values based on PD and FD coincide.

#### Proposition 3

If 𝔽 = *F*[*T, h*], then *φ*_𝔽_(*x*) is equal to the Fair Proportion index for taxon *x* on tree *T* for the edge length assignment ℓ_*h*_.

*Proof*. Let 𝔽 = *F*[*T, h*] and let *x* ∈ *X*. As noted above, we have:

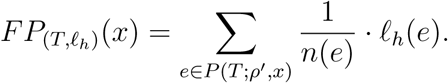

Importantly, we can also write *φ*_𝔽_(*x*) as follows:

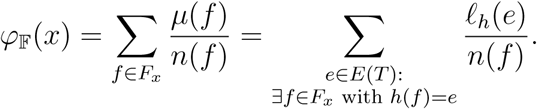

Now, since 𝔽 = *F*[*T, h*], all the edges on which features present in *F*_*x*_ arise must lie on the unique path from *ρ*′ to *x*. Moreover, a feature *f* ′ not contained in *F*_*x*_, cannot have arisen on this path. More precisely, if a feature *f* arises on edge *e*, then a taxon *x* ∈ *X* has this feature if and only if it is a descendant of *e*. In particular, *n*(*f*) = *n*(*e*). In summary, this implies that 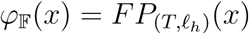. □

We now establish a further result. We show that *φ*_𝔽_(*x*) can always be interpreted as *FP*_(*T*,ℓ)_(*x*) for any tree *T* (even if 𝔽 ≠ *F*[*T* ′, *h*] for any tree *T* ′).

#### Theorem 2

Let 𝔽 be a feature assignment such that *F*_*x*_ ≠ ø for all *x* ∈ *X*, and let *T* be any rooted phylogenetic *X*-tree (with additional stem edge). Then, there exists an edge length assignment ℓ : *E* → ℝ^>0^ that assigns strictly positive lengths to all edges of *T*, such that *φ*_𝔽_(*x*) = *FP*_(*T*,ℓ)_(*x*) for all *x* ∈ *X*.

The proof of Theorem 2 is provided in the Appendix, however we provide an outline of the argument here. First observe that if *T* is a star tree then Theorem 2 clearly holds, since we can simply assign edge length *φ*_𝔽_(*x*) to the edge incident with leaf *x* and obtain *φ*_𝔽_(*x*) = *FP*_(*T*,ℓ)_(*x*) for all *x* ∈ *X*. If *T* is not a star tree, then we could assign edge length 0 to all the interior edges and the stem edge, and apply the same trick to obtain the required identity. The non-trivial part of the proof of Theorem 2 is to show that one can ‘lift’ some fraction of the lengths of the pendant edges so as that (i) *all* edges of *T* have strictly positive length, and in such a way that (ii) the required identity between the FD and PD diversity indices holds for each taxon *x*.

## Discussion

It would be a mistake to interpret Theorem 1 above as stating that feature diversity coincides with phylogenetic diversity (on a given tree with suitably chosen branch lengths) only under evolutionary scenarios in which features arise once in the tree and are never lost. Instead, Theorem 1 states that these two measures coincide precisely when the distribution of features across taxa can be described by such a single-gain-and-no-loss model, even if the underlying reality might be different. For instance, a feature can arise along a stem edge, be lost in one of the two descendant edges, but arise again in its descendants such that the entire crown clade expresses the feature. The feature’s true history is obscured but its distribution is still perfectly congruent with the underlying tree and thus meets the conditions of Proposition 1.

This points to one scenario where Theorem 1 will not hold, where the rate of evolution is high enough and the state space is small enough (e.g., features are discrete and can be both gained and lost) that at least some features have non-trivial probabilities of arising more than once on a tree. Faith (1992) astutely pointed out that such convergent features “are not predictive of similarities of other features,” such that “greater phylogenetic diversity will, on average, imply greater feature diversity as defined by any particular collection of features.” Because empirical measures of PD and measured FD need not coincide (Devictor et al. (2010); see also the discussion in Winter et al. (2013)), one critical question is whether there are subsets of features that simultaneously (i) are more valuable to conservation than the average feature and (ii) are (or are likely to be) convergent, perhaps due to parallel adaptation (Mazel et al. (2018, 2019); Owen et al. (2019)). To the extent that there are, the force of Faith’s all-important “average” PD = FD statement weakens. However, answering the question is non-trivial because it requires that we know about the mode of evolution of conservation-relevant features in a focal clade. The only attempt to test this we know of is by Forest et al. (2007) for Southern African plants, where the patterns supported Faith’s average argument. More tests would be welcome.

A second (related) reason why Theorem 1 allows PD and FD to diverge in applications is that even when 𝔽 = *F*[*T, h*], the edge lengths must be suitably chosen. Under a stochastic process in which features arise independently at a constant (and very small) rate *r*, then conditional on a feature arising (at least) once in the tree, as *r* → 0, the expected number of features that arise on an edge will be proportional to the temporal length of that edge (and each trait will arise exactly once in the tree). This (coupled with Faith’s “average” argument) is the justification for using time-calibrated ultrametric phylogenetic trees when comparing PD scores. However, the evolution of some important subset of features may not be captured with this model at all (Mazel et al. (2017)), or, more prosaically, may simply evolve at such a high rate that the time-calibrated ultrametric tree edge lengths are not predictive of the number and placement of features (for example, due to saturation). Here again, both theoretical (see e.g. Tucker et al. (2018)) and empirical tests using features of known conservation value are needed.

Two observations can be made on the basis of Theorem 2. The first is that the compact expression for the *φ*_𝔽_(*x*) measure of feature diversity does not require any particular model of feature evolution on a tree: different features and different subtrees can be governed by different processes. While we might still require Faith (1992)’s “average” argument, namely, that the distribution of measured features mirrors the features of conservation concern more generally, this might expand its usefulness. Once again, this is open to empirical testing. However, the Shapley value is quite specific in what it measures: it is the expected contribution (here, of features) to all possible future subsets of taxa, where subset sizes are equiprobable (Steel, 2016). This is a strong assumption that bears further scrutiny (see also Faith (2008)).

In conclusion, our paper provides a precise mathematical framework to help address some fundamental questions and possible future approaches concerning the link between feature and phylogenetic diversity, a critical connection for phylogeny-oriented conservation triage.

## Proofs of Propositions

### Proof of Proposition 1

i. First, suppose that 𝔽 = *F*[*T, h*], and let *f* ∈ ℱ be an arbitrary feature. Moreover, let *e* = *h*(*f*) be the edge *f* arose on. Then, precisely those leaves of *T* that are descendants of *e* contain feature *f*. In particular, *X*_*f*_ corresponds to a cluster of *T* (namely, to the cluster of leaves descending from *e*). Now, suppose that *X*_*f*_ corresponds to a cluster of *T* for each feature *f* ∈ ℱ. Then we can realize 𝔽 on *T* by setting *h*(*f*) to the edge directly above the most recent common ancestor of the taxa in *X*_*f*_ for each *f* ∈ ℱ. In particular, 𝔽 = *F*[*T, h*]. Moreover, when 𝔽 = *F*[*T, h*], *h*(*f*) must be the edge, say *e*, directly above the most recent common ancestor of the taxa in *X*_*f*_. If *f* had arisen above *e*, say on some edge *e*′, there would be at least one taxon *x* in the cluster *c*_*T*_ (*e*′) induced by *e*′ (i.e. in the set of leaves of *T* that are separated from the root of *T* by *e*′) with *x* ∈ *c*_*T*_ (*e*′) \ *X*_*f*_. This, however, would imply that *f* was lost on the way from *e*′ to *x*, which is a contradiction. Similarly, if *f* had arisen below edge *e*, say on some edge *e*″, this would imply that the cluster *c*_*T*_ (*e*″) induced by *e*″ was a strict subset of *X*_*f*_, i.e. *c*_*T*_ (*e*″) ⊂ *X*_*f*_. In other words, there would be at least one taxon *y* ∈ *X*_*f*_ \ *c*_*T*_ (*e*″), which implies that *f* arose at least twice in *T*. This is again a contradiction.
ii. First, suppose that 𝔽 = *F*[*T, h*]. By Part (i) of this Lemma, this implies that the set *X*_*f*_ corresponds to a cluster of *T* for all *f* ∈ ℱ. In particular, 𝒞_ℱ_ = {*X*_*f*_ : *f* ∈ ℱ} is a set of clusters induced by a rooted phylogenetic *X*-tree *T*, which implies that 𝒞_ℱ_ is a hierarchy on *X* (cf. Proposition 2.1 in Steel (2016)). Now, suppose that 𝒞_ℱ_ is a hierarchy on *X*. This implies (again by Proposition 2.1 in Steel (2016)) that 𝒞_ℱ_ is the set of clusters of some rooted phylogenetic *X*-tree *T*. By Part (i) of this Lemma, this in turn implies that 𝔽 = *F*[*T, h*].□

### Proof of Proposition 2

Notice that both *φ*_𝔽_(*x*) and *ν*_𝔽_(*Y*) (with *Y* ⊆ *X*) are linear functions in *µ*(*f*). More precisely,

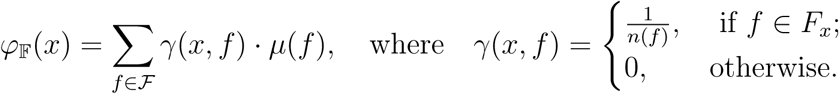

Analogously,

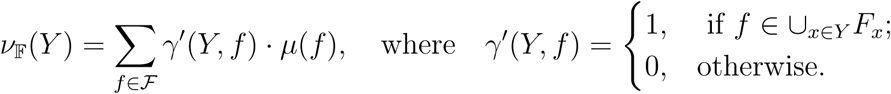

Thus, by linearity (see also Lemma 6.14 in Steel (2016)), it suffices to show the statements for the case that one element of ℱ, say *f*_*i*_, has score *µ*(*f*_*i*_) = 1, whereas *µ*(*f*_*j*_) = 0 for all *f*_*j*_ ∈ ℱ \ {*f*_*i*_} (note that *µ* was earlier defined to be strictly positive, but we are here relaxing this for algebraic convenience).

For the first part of the proof, recall that given the cooperative game (*X, ν*_𝔽_), the Shapley value of *x* ∈ *X* is defined as (Shapley (1953))

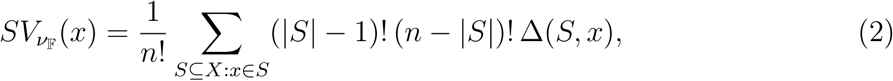

where Δ(*S, x*) = *ν*_𝔽_(*S*) − *ν*_𝔽_(*S* \ {*i*}). We now show that 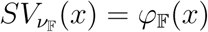 (where we assume that *µ*(*f*_*i*_) = 1 and *µ*(*f*_*j*_) = 0 for all *f*_*j*_ ∈ ℱ \ {*f*_*i*_}).

We can distinguish two cases:

- If *f*_*i*_ ∉ *F*_*x*_, then Δ(*S, x*) = 0 for all *S*, and thus, 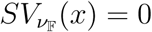. On the other hand, we clearly also have *φ*_𝔽_(*x*) = 0.
- If *f*_*i*_ ∈ *F*_*x*_, then Δ(*S, x*) = 1 if (i) *x* ∈ *S* and (ii) there is no *y* ∈ *S* with *f*_*i*_ ∈ *F*_*y*_; otherwise Δ(*S, x*) = 0. Let *C* ⊆ *X* be the set of taxa that have feature *f*_*i*_, i.e., *C* = {*y* ∈ *X* : *f*_*i*_ ∈ *F*_*y*_}, and so *n*(*f*_*i*_) = |*C*|. Then, 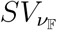 can be written as

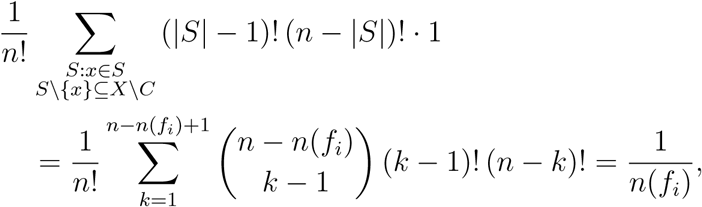

where the last equality follows from the fact that 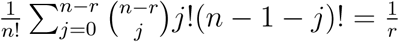 for 1 ⩽ *r* ⩽ *n* (Lemma 6.15 in Steel (2016)) (here: *j* = *k* − 1 and *r* = *n*(*f*_*i*_)). On the other hand, 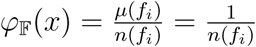, which completes the proof.

The second part of Proposition 2, follows directly from properties of the Shapley value; however we give a direct proof. Again, it suffices to consider the case where *µ*(*f*_*i*_) = 1 and *µ*(*f*_*j*_) = 0, for all *j* ≠ *i*, in which case we obtain the required equality:

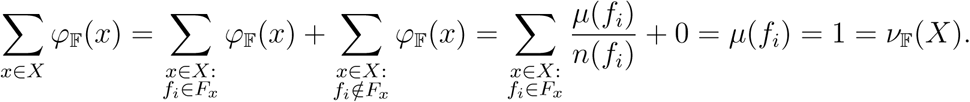

□

### Proof of Inequality (1).

Let *W*(*Y*) := ⋃_*x*∈*Y*_ *F*_*x*_. From the proof of Proposition 2 we have:

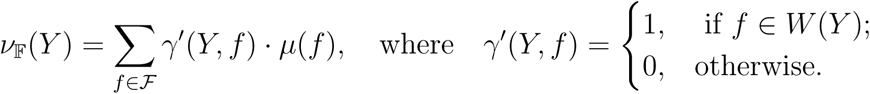

Now,

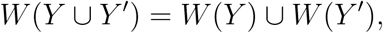

and

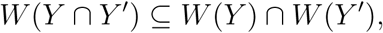

(and the containment can be strict). It follows that for all *f* ∈ ℱ and all *Y, Y* ′ ⊆ *X*.:

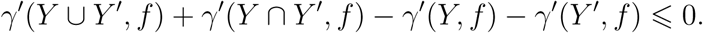

Since *ν*_𝔽_(*Y ∪ Y*) + *ν*_𝔽_(*Y ∩ Y* ′) − *ν*_𝔽_(*Y*) − *ν*_𝔽_(*Y* ′) is a positive weighted sum of the corresponding *γ*′ quantities above, Inequality (1) now follows.□

## Acknowledgements

MS thanks the Royal Society of NZ (Catalyst Grant) for funding support. AOM is supported by the Natural Sciences and Engineering Research Council of Canada, and is grateful to members of the iDiv synthesis centre working group “Conservation and Phylogenies” for ongoing discussion, and Dan Faith for continual engagement on this topic.

## Appendix: Proof of Lemmas 1–3, and Theorem 2

### Proof of Lemma 1.

Suppose that 𝔽 = *F*[*T, h*]. For each *f* ∈ ℱ, let *X*_*f*_ be the set of taxa that have feature *f*. Then,

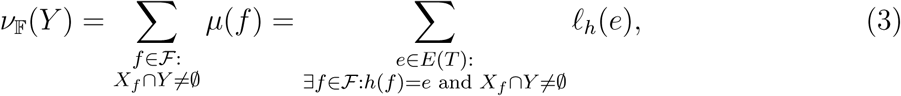

where the last equality follows from the fact that 𝔽 = *F*[*T, h*].

On the other hand,

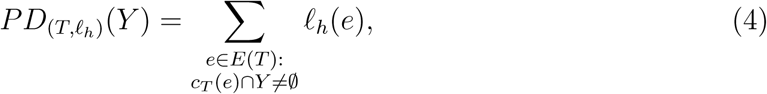

where *c*_*T*_ (*e*) denotes the set of leaves of *T* that are separated from the root of *T* by *e*. Now, as 𝔽 = *F*[*T, h*], when *e* = *h*(*f*), *c*_*T*_ (*e*) corresponds to the set *X*_*f*_. Thus, for *e* ∈ *E*(*T*), we can conclude that *c*_*T*_ (*e*) *∩ Y* ≠ ø precisely if

- ∃*f* ∈ ℱ : *h*(*f*) = *e* and *X*_*f*_ *∩ Y* ≠ ø, or
- ∄*f* ∈ ℱ : *h*(*f*) = *e* and *e* is a pendant edge incident to a leaf *y* ∈ *Y* (in which case ℓ_*h*_(*e*) = 0).

Thus, we can re-write Eqn. (4) as

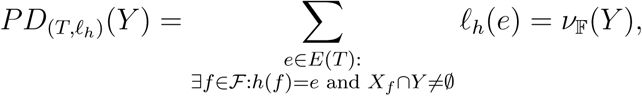

where the last equality follows from Eqn. (3). This completes the proof. □

### Proof of Lemma 2:.

We will prove this statement by contradiction. Thus, assume that *ν*_𝔽_(*Y*) = *PD*_(*T*,ℓ)_(*Y*) for all *Y* ⊆ *X* but there is no map *h* : ℱ → *E* such that 𝔽 = *F*[*T, h*]. We now distinguish two cases: (i) 𝔽 cannot be explained by *T*, but by some other tree *T* ′, i.e. 𝔽 = *F*[*T* ′, *h*′], or (ii) the collection of sets 𝒞_ℱ_ = {*X*_*f*_ : *f* ∈ ℱ} does not form a hierarchy and cannot be explained by any tree (cf. Proposition 1, Part (ii)).

i. First, suppose that 𝔽 ≠ *F*[*T, h*] but 𝔽 = *F*[*T* ′, *h*′], and *ν*_𝔽_(*Y*) = *PD*_(*T*,ℓ)_(*Y*) for all *Y* ⊆ *X*. Now, as *T* ≠ *T*′, there must be some *i, j, k* ∈ *X* such that restricting *T* and *T* ′ to {*i, j, k*} yields distinct trees. More precisely, there exist *i, j, k* ∈ *X* such that
  a. *T* |_{*i,j,k*}_ is either the caterpillar tree on three leaves with cherry [*i, j*] or the rooted star tree on {*i, j, k*},
  b. *T* ′|_{*i,j,k*}_ is either the caterpillar tree on three leaves with cherry [*i, k*] or the rooted star tree on {*i, j, k*},
  c. *T* |_{*i,j,k*}_ ≠ *T* ′|_{*i,j,k*}_ (in particular, *T* |_{*i,j,k*}_ and *T* ′|_{*i,j,k*}_ are not both star trees).
  Let Δ_𝔽_(*x, x*′) := *ν*_𝔽_({*x*}) + *ν*_𝔽_({*x*′}) − *ν*_𝔽_({*x, x*′}), for each distinct pair *x, x*′ ∈ *X*. Then as *ν*_𝔽_(*Y*) = *PD*_(*T*,ℓ)_ for all *Y* ⊆ *X*, we have from (a) that:

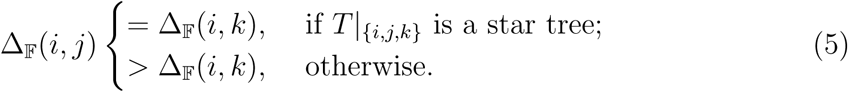 On the other hand, as 𝔽 = *F*[*T* ′, *h*′], we have by Lemma 1, that 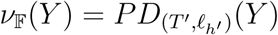 for all *Y* ⊆ *X* (where ℓ_*h*′_ (*e*) = ∑_*f : h′(f)=e*_ *µ*(*f*); in particular, ℓ_*h*′_ (*e*) > 0 for each interior edge *e* of *T* ′). This implies that:

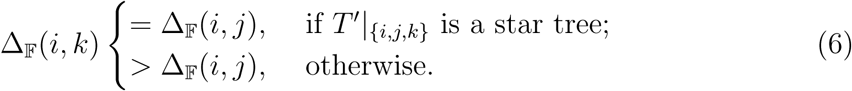 Comparing Eqns. (5) and (6), and using the fact that *T* |_{*i,j,k*}_ and *T* ′|_{*i,j,k*}_ cannot both be star trees, this yields a contradiction. As (*i, j, k*) was an arbitrary triple of leaves for which *T* |_{*i,j,k*}_ *T* ′|_{*i,j,k*}_, this contradiction implies that the initial assumption was wrong. In particular, 𝔽 = *F*[*T, h*].
ii. Now, assume that *ν*_𝔽_(*Y*) = *PD*_(*T*,ℓ)_(*Y*) for all *Y* ⊆ *X*, but 𝒞_ℱ_ = {*X*_*f*_ : *f* ∈ ℱ} does not form a hierarchy. This implies that there exists *f*_1_, *f*_2_ ∈ ℱ such that
  a. There exists a taxon *x*_1_ ∈ *X* such that 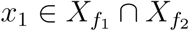.
  b. There exists a taxon *x*_2_ ∈ *X* such that 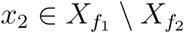.
  c. There exists a taxon *x*_3_ ∈ *X* such that 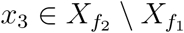.

We now partition the feature set ℱ into eight pairwise disjoint subsets *A*,…, *G*, where

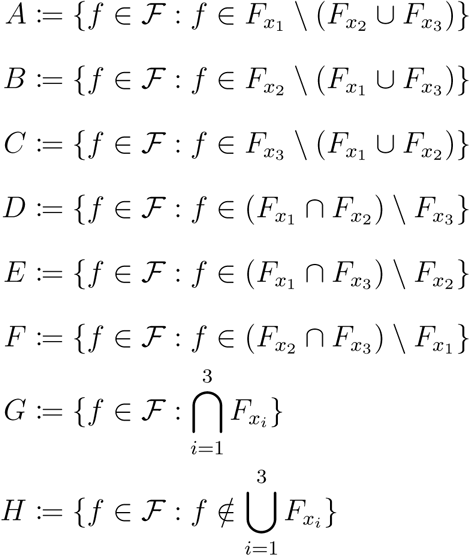

Note that *D* ≠ ø (because by 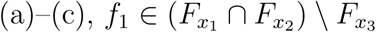). Analogously, *E* ≠ ø (because 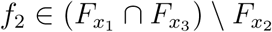).

Given a set of features *S*, let *µ*(*S*) := ∑_*f* ∈*S*_ *µ*(*f*) denote the sum of scores of features present in *S*. As *µ*(*f*) > 0 for all *f* ∈ ℱ, by the preceding argument, in particular *µ*(*D*), *µ*(*E*) > 0.

We now compute *ν*_𝔽_(*Y*) for all *Y* ⊆ {*x*_1_, *x*_2_, *x*_3_} with |*Y*| ⩾ 1, and compare it to *PD*_(*T*,ℓ)_(*Y*). Recall that *PD*_(*T*,ℓ)_(*Y*) for *Y* ⊆ *X* is computed by considering the sum of edge lengths in the minimum subtree of *T* connecting the taxa in *Y* and *ρ*′. Without loss of generality, we can assume that the subtree induced by {*x*_1_, *x*_2_, *x*_2_} has the structure depicted in Fig. 2 (otherwise, we exchange leaf labels).

Now, by assumption *ν*_𝔽_(*Y*) = *PD*_(*T*,ℓ)_(*Y*) for all *Y* ⊆ *X*. For *Y* ⊆ {*x*_1_, *x*_2_, *x*_3_} with |*Y* | ⩾ 1, this gives rise to a system of 7 linear equations (where ℓ(*p*) denotes the length of path *p*):

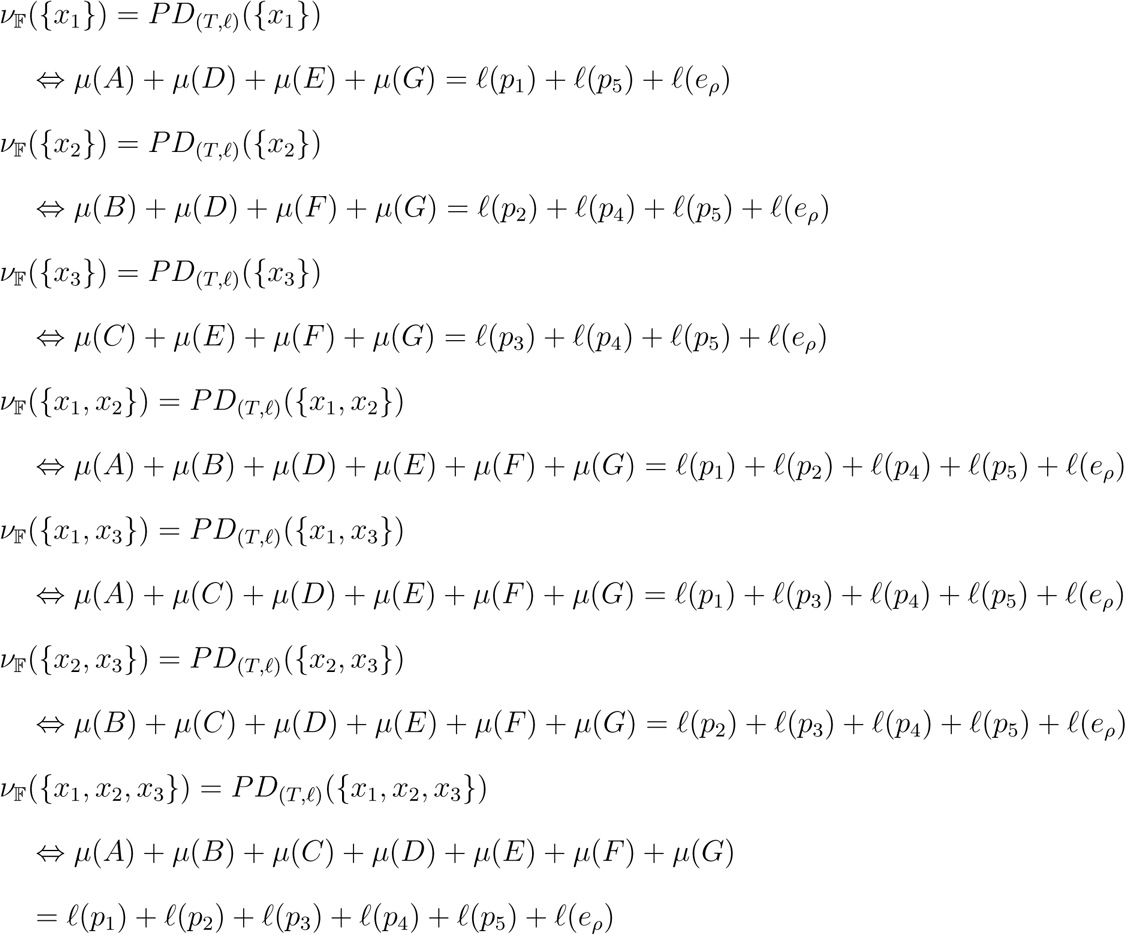

Solving this system of linear equations for *µ*(*A*),…, *µ*(*G*) yields *µ*(*A*) = ℓ(*p*_1_), *µ*(*B*) = ℓ(*p*_2_), *µ*(*C*) = ℓ(*p*_3_), *µ*(*D*) = *µ*(*E*) = 0, *µ*(*F*) = ℓ(*p*_4_), and *µ*(*G*) = ℓ(*p*_5_) + ℓ(*e*_*ρ*_).

However, as our assumption implies that *µ*(*D*), *µ*(*E*) > 0, this is a contradiction. Thus, the initial assumption was false. In particular, {*X*_*f*_ : *f* ∈ ℱ} forms a hierarchy. Thus, by Proposition 1, Part (ii), there exist *T* ′ and *h*′ such that 𝔽 = *F*[*T* ′, *h*′]. Now, by case (i) of this proof, this implies 𝔽 = *F*[*T, h*]. This completes the proof.□

### Proof of Lemma 3:.

Let *T* be a rooted phylogenetic *X*-tree (with additional stem edge), and assume that *PD*_*T*_ (*Y*) is given for all *Y* ⊆ *X* with |*Y* | ⩽ 2. We now show that we can uniquely infer the edge lengths of *T* from these scores. Let *i* ∈ *X* be a leaf of *T*. Then, there is a unique path *e*_*k*+1_, *e*_*k*_,…, *e*_1_, *e*_0_ from *ρ*′ to *i* in *T* (see Fig. 1), and we can infer the lengths of these edges in a ‘top-down’ approach (i.e., starting with edge *e*_*k*+1_ and moving down the tree towards edge *e*_0_).

For ℓ(*e*_*k*+1_), let *j* be a leaf that is not a descendant of edge *e*_*k*_ (in other words, *j* is not in the same maximal pending subtree as *i*). Then, clearly,

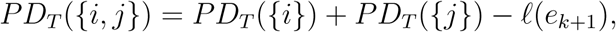

(because ℓ(*e*_*k*+1_) contributes twice to the sum *PD*_*T*_ ({*i*}) + *PD*_*T*_ ({*j*}), but only once to *PD*_*T*_ ({*i, j*})). In other words, ℓ(*e*_*k*+1_) = *PD*_*T*_ ({*i*}) + *PD*_*T*_ ({*j*}) − *PD*_*T*_ ({*i, j*}).

Now, let *e*_*i*_ = (*u, v*) be an interior edge in the path from *ρ*′ to *i*, for which the lengths of its preceding edges are already determined, i.e., ℓ(*e*_*k*+1_),…, ℓ(*e*_*i*+1_) are known. Moreover, let *j* be a leaf that is a descendant from *e*_*i*_, but not from *e*_*i*−1_.

Then, with a similar argument as in the previous case, we have

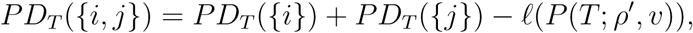

where ℓ(*P* (*T*; *ρ*′, *v*)) denotes the length of the unique path from *ρ*′ to *v* in *T* (which contributes twice to the sum *PD*_*T*_ ({*i*}) + *PD*_*T*_ ({*j*}), but only once to *PD*_*T*_ ({*i, j*})). In other words, ℓ(*P* (*T*; *ρ*′, *v*)) = *PD*_*T*_ ({*i*}) + *PD*_*T*_ ({*j*}) − *PD*_*T*_ ({*i, j*}). On the other hand, ℓ(*P* (*T*; *ρ*′, *v*)) = ℓ(*e*_*k*+1_) + ℓ(*e*_*k*_) +… + ℓ(*e*_*i*+1_) + ℓ(*e*_*i*_), and as ℓ(*e*_*k*+1_),…, ℓ(*e*_*i*+1_) are known, we can uniquely infer ℓ(*e*_*i*_).

Finally, after inferring the lengths of the edges *e*_*k*+1_, *e*_*k*_,…, *e*_1_ as described above, we can also uniquely infer the length of the pendant edge *e*_0_ incident to *i* as 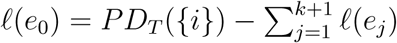.

In summary, we can uniquely infer all edge lengths of edges in the path from *ρ*′ to *I* from the PD scores of subsets of *X* of size at most 2. As *i* was an arbitrary leaf of *T*, this completes the proof. □

### Proof of Theorem 2.

Let 𝔽 be a feature assignment and let *T* be a rooted phylogenetic *X*-tree. First, note that we can always achieve *φ*_𝔽_(*x*) = *FP*_(*T*,ℓ′_)(*x*) for all *x* ∈ *X* when we consider an edge length assignment ℓ′ that allows edges to be assigned length zero because, in this case, if *e*_*x*_ denotes the pendant edge incident to *x*, we can set ℓ′(*e*_*x*_) = *φ*_𝔽_(*x*) for each *x* ∈ *X*, and ℓ′(*e*) = 0 for all interior edges and the stem edge, which clearly results in *φ*_𝔽_(*x*) = *FP*_(*T*,ℓ′_)(*x*) for all *x* ∈ *X*.

We now show that we can obtain an edge length assignment ℓ assigning strictly positive lengths to all edges of *T* from ℓ′ by redistributing lengths in a ‘bottom-up’ approach (i.e. moving from pendant edges towards the stem edge).

First, for each pendant edge *e*_*x*_, set ℓ(*e*_*x*_) = ℓ′(*e*_*x*_), which is strictly positive, due to the assumed condition *F*_*x*_ ≠ ø, along with the fact that *µ* takes strictly positive values. Now, let *e* be an edge of *T* such that all edges descending from *e* already have strictly positive lengths, whereas all edges above *e* (if they exist) still have length zero. Let *e*_1_,…, *e*_*k*_ denote the descending edges incident to *e*, and let *t*_1_,…, *t*_*k*_ denote the subtrees pending from *e* (where tree *t*_*i*_ has stem edge *e*_*i*_ for *i* = 1,…, *k*). Moreover, for *i* = 1,…, *k*, let 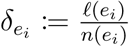 denote the ratio between the length of *e*_*i*_ and the number of leaves descending from it. Without loss of generality, we may assume that edge *e*_1_ minimizes this ratio (else we exchange edge labels). Furthermore, let 0 < *c* < 1. We now re-assign edge lengths to *e*_1_,…, *e*_*k*_ and *e* as follows (where ℓ_old_(*e*_*i*_) refers to the edge length *e*_*i*_ is currently assigned):

1. 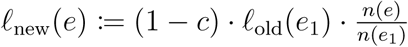
2. ℓ_new_(*e*_1_) := *c* · ℓ_old_(*e*_1_),
3. 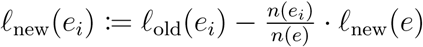 for *i* = 2, …, *k*.

Now, in order to show that this is a valid re-distribution of edge lengths, we need to show that

i. ℓ_*new*_(*e*) > 0 and ℓ_new_(*e*_*i*_) > 0 for *i* = 1,…, *k*.
ii. 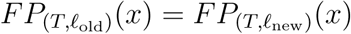 for all *x* ∈ *X*.

First, consider (i). As ℓ_old_(*e*_1_) > 0 by assumption, and 0 < *c* < 1, we clearly have ℓ_new_(*e*) > 0, and ℓ_new_(*e*_1_) > 0. Now, consider *e*_*i*_ for *i* ∈ {2,…, *k*}. Here, we have

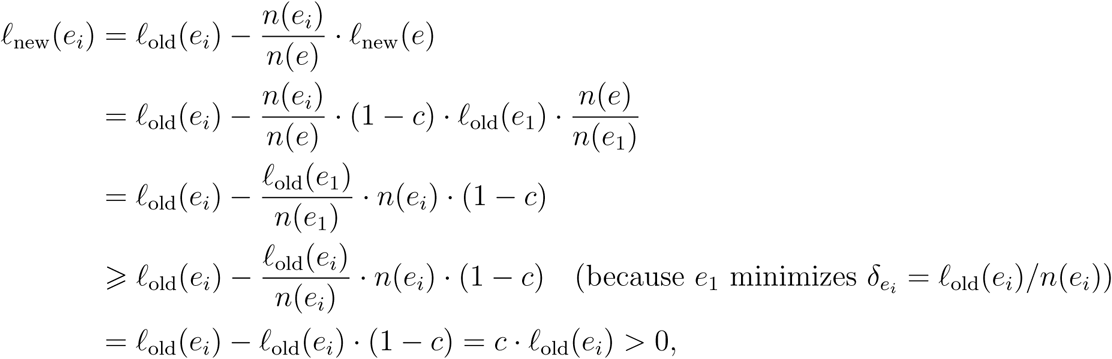

where the last inequality again follows from the fact that (by assumption) ℓ_old_(*e*_*i*_) > 0 and 0 < *c* < 1. This completes the proof of (i).

For (ii) note that the FP indices of taxa not descending from *e* are not affected by the re-assignment of edge lengths, so it suffices to consider all *x* ∈ *c*_*T*_ (*e*). In the following, let *t*_*i*_ \ *e*_*i*_ be the rooted phylogenetic tree obtained from *t*_*i*_ by deleting its stem edge. Then, we clearly have for all *x* ∈ *c*_*T*_ (*e*):

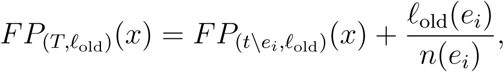

(because by assumption all edges above *e*_*i*_ have length zero before the re-assignment of edge lengths according to steps 1–3). On the other hand, we have for all *x* ∈ *c*_*T*_ (*e*):

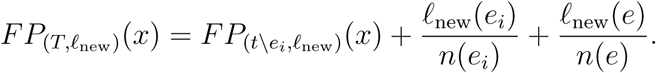

Note that 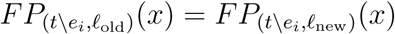 for all *x* ∈ *c*_*T*_ (*e*) (because the lengths of edges in *t*_*i*_ \ *e*_*i*_ are not changed). We now show that 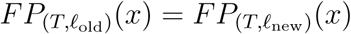 for all *x* ∈ *X*. First, let *x* ∈ *t*_1_. Then, we have

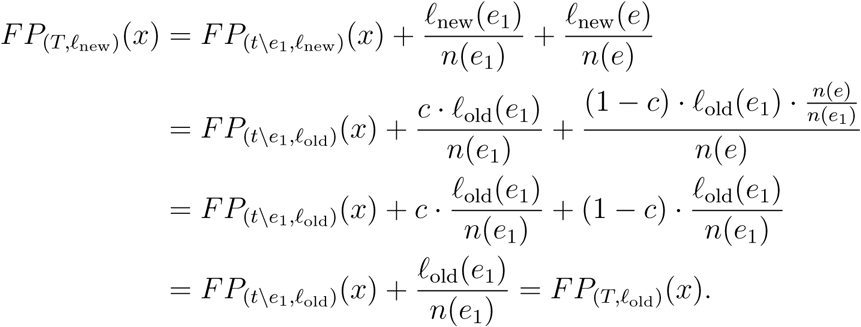

Now, let *x* ∈ *t*_*i*_ for *i* ∈ {2,…, *k*}. Then, we have

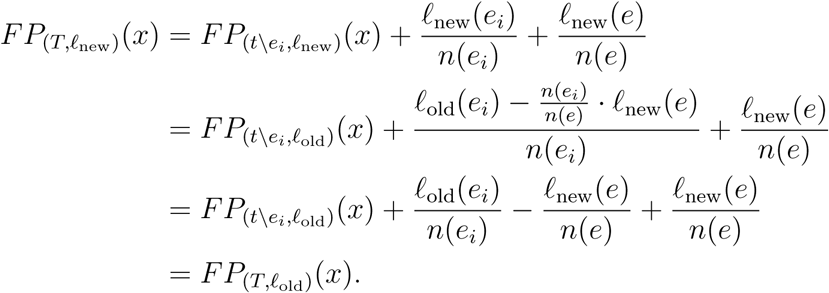

In summary, re-assigning edge lengths according to the conditions 1–3 (listed above) is valid (because conditions (i) and (ii) hold). Thus, for each edge *e* whose length was changed, we now simply set ℓ(*e*) = ℓ_new_(*e*) and repeat the procedure. In this way, we can construct an edge length assignment ℓ that assigns strictly positive lengths to *all* edges of *T* (including pendant edges and the stem edge), such that *φ*_𝔽_(*x*) = *FP*_(*T*,ℓ)_(*x*) for all *x* ∈ *X*. This completes the proof. □

